# Systemic lipid peroxidation and skeletal tissues metabolism in animals with simulated posttraumatic osteoarthrosis

**DOI:** 10.1101/2021.11.30.470525

**Authors:** Svetlana V. Belova, Roman A. Zubavlenko, Vladimir Yu. Ulyanov

## Abstract

The role of individual factors in the progressing of degenerative and dystrophic changes of articular cartilage as the focus of pathologic changes in posttraumatic osteoarthrosis (PTOA) has been studied comprehensively. However, no single concept of pathogenetic mechanisms leading to the formation of permanent changes in the affected joint has been represented yet. Therefore, the investigation of lipid peroxidation and skeletal tissues metabolism in animals with simulated PTOA is a challenging and important issue.

In 28 non-linear male rats (12 intact control animals and 16 experimental animals with simulated knee OA) we studied the lipid peroxidation rate by the level of lipid hydroperoxides, antioxidant system activity by the indicants of total antioxidant, and thiol statuses. The destruction processes in skeletal tissues were evaluated by the level of hyaluronan, a cartilaginous tissue marker, and subchondral bone turnover makers (fibroblast growth factor-23, osteoprotegerin, sclerostin, osteocalcin).

In rats with simulated PTOA, we observed the activation of lipid peroxidation with the accumulation of lipid hydroperoxides in systemic blood flow along with the insignificant decrease of thiol status indicant while the normal antioxidant activity is maintained. We also revealed the significant increase of biopolymer hyaluronan (p<0.05) in a negative change of turnover regulation marker (the significant increase in fibroblast growth factor-23 (p<0.05) with the emerging trend for osteoprotegerin and sclerostin decrease) and bone formation marker (high osteocalcin) as compared to the intact controls.

Our findings confirmed that animals with simulated PTOA featured rearranging of skeletal tissues as compared to intact animals. This process involved degenerative destruction of both chondral and osseous tissues along with profound prooxidant activity and relative incompetence of the thiol system while the indicants of general antioxidant activity remain normal. This suggests the importance of the analyzed markers in PTOA pathogenesis and determined the prospects of their use for the evaluation of therapy efficacy.

## Background

Posttraumatic osteoarthrosis (PTOA) results from traumatic changes in the joints and affects a large number of active age individuals [Golovach I.Yu. Yeghudina Ye.D., 2019]. The activation of lipid peroxidation in PTOA is known to produce toxic metabolites destructing the joint components [Zahvatov A.N., et al., 2018] that contain skeletal tissues including chondral and osseous tissues performing mechanical and metabolic functions in the body.

In PTOA the role of individual factors in the emergence of degenerative and dystrophic changes in articular cartilage as the focus of pathology changes has been studied comprehensively. However, no single concept of pathogenetic mechanisms leading to the formation of permanent changes in the affected joint has been represented yet.

This research had the evaluation of systemic lipid peroxidation and skeletal tissues metabolism in animals with simulated PTOA is a challenging and important issue as its **objective**.

## Material and Methods

The research involved 28 non-linear male rats weighing 270-310 g divided into two groups: 12 intact animals made up the control group and 16 animals with simulated knee PTOA made up the experimental group.

### Entry criteria

1. Male rats
2. 250-350 g of body mass
3. 3 to 6 months of age
4. No signs of other diseases

## Research Design

This experimental study was performed following the European Convention for the Protection of Vertebrate Animals used for Experimental and Other Scientific Purposes (Strasbourg, 1986), and the Russian Federation Ministry of Healthcare Order #267 of June 19, 2003, On approval of laboratory practice regulations.

Knee PTOA was simulated by anterior cruciate ligament transections in intercondylar space with the damage to cartilage and subchondral bones caused by the tip of the scalpel that formed the wedge-shaped defects on the surface of femoral articular cartilage followed by layered closure of the wounds [Slesarenko N.A., Shirokova E.O., 2012].

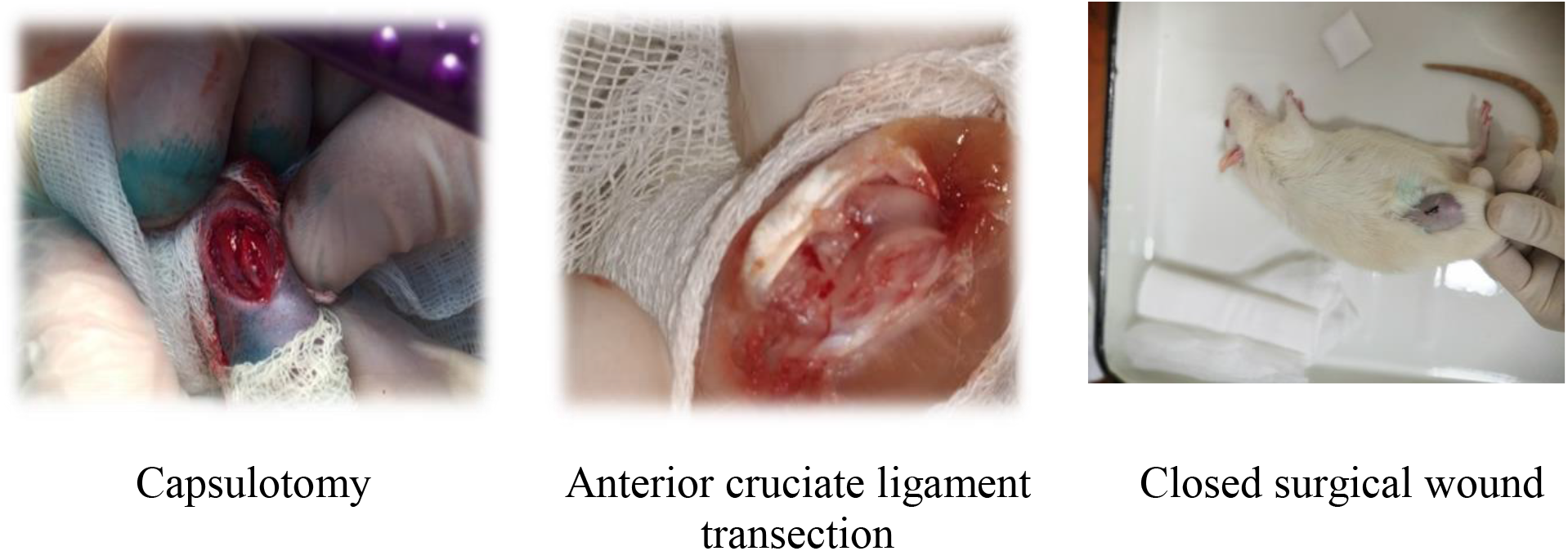

- The rate of systemic lipid peroxidation was determined by the serum levels of lipid hydroperoxides. The activity of the antioxidant system was evaluated by the indicants of total antioxidant and thiolic statuses.
- The state of skeletal tissues was evaluated by cartilaginous tissue marker (hyaluronan) and subchondral bone turnover makers (fibroblast growth factor-23, osteoprotegerin, sclerostin, osteocalcin).

Blood samples to perform tests were taken by intracardiac collection from anesthetized animals. The findings were statistically processed in Microsoft Excel 2010 using the Mann-Whitney U-test; the results were presented as a median [Me] and interquartiles [Q (25; 75)]. The differences were considered significant in p<0.05.

## Results

By Day 28 of the experiment, we observed the rise of local temperature in the knee areas and limitation of experimental joint movement in experimental rats as compared to the initial values.

The analysis of systemic lipid peroxidation progress revealed its activation with the accumulation of lipid hydroperoxides (see Table 1). These intermediate molecular products generally destabilize plasmatic membranes [Martusevich A.A., et al, 2012; Yusupov M., et. al., 2017].

**Table 1.** Indicators of systemic lipid peroxidation and antioxidant system activity.

| Indicators | Controls | Experimental |
| --- | --- | --- |
| Lipid hydroperoxides (nmol) | 6.3 (5.7; 7.3)<br>n=10 | 7.6 (6,9; 9.7)<br>p=0.03027*<br>n=14 |
| Total antioxidant status<br>($\mu\text{mol/l}$) | 302.3 (280.1; 317.1)<br>n=11 | 285.4 (272.5; 311.8)<br>p=0.31117<br>n=14 |
| Thiol status ($\mu\text{mol/l}$) | 108.3 (102.3; 117.4)<br>n=12 | 90.8 (78.3; 105.8)<br>p=0.01677*<br>n=14 |

*Note:* p – difference between the controls and experimental group; \* – significant difference; n – number of animals. The findings are presented as a median [Me] and interquartiles [Q (25; 75)].

The analysis of the general state of the antioxidant support network revealed no significant differences between the analyzed groups of animals. However, in the rats with PTOA we observed a significant decrease in thiol compound activity (see Table 1) possibly suggesting the first signs of stress in the redox-buffer cell system.

The profound prooxidant activity caused by the intensification of the reactive *oxygen* intermediate formation along with the incompetence of individual links in the antioxidant system in PTOA animals was associated with degeneration and destruction of extracellular matrix evidenced by the significant increase of serum hyaluronan (see Table 2). This was probably caused by its depolymerization in unfavorable metabolic conditions [Zahvatov A.N., et al, 2018; Shimura Y. et al., 2018].

**Table 2.** Skeletal tissue response markers.

| Markers | Controls | Experimental |
| --- | --- | --- |
| Hyaluronan (ng/ml) | 62.6 (58.2; 66.2)<br>n=10 | 101.7 (91.4; 108.5)<br>p=0.0000318*<br>n=15 |
| Osteoprotegerin (pg/ml) | 1177.9 (981.3; 1620.1)<br>n=12 | 1010.1 (937.2; 1090.1)<br>p=0.125526<br>n=16 |
| Fibroblast growth factor-23 (pg/ml) | 233.6 (201.5; 248.6)<br>(n=12) | 278.4 (262.6; 337.8)<br>p=0.000763*<br>n=16 |
| Sclerostin (pg/ml) | 206.5 (147.7; 244.3)<br>n=12 | 169.6 (146.1; 235.3)<br>p=0.710347 |
|  |  | n=16 |
| Osteocalcin (ng/ml) | 189.8 (146.6; 211.0)<br>n=12 | 209.1 (178.2; 141.6)<br>p=0.110831<br>n=14 |

*Note:* p – difference between the controls and experimental group; \* – significant difference; n – number of animals. The findings are presented as a median [Me] and interquartiles [Q (25; 75)].

The analysis of subchondral bone remodeling in PTOA animals revealed a significant rise (p<0.05) in fibroblast growth factor-23 as compared to the controls. AΠA-23 is a phosphaturic hormone expressed in osseous tissue and synthesized by osteocytes, that controls phosphate homeostasis and vitamin D [Kuznik B.I., et al, 2017]. We also observed the increase (p>0.05) in concentrations of osteocalcin (see Table 2), a marker of bone formation, suggesting disorder in osseous tissue remodeling. Sclerostin and osteoprotegerin, the makers of bone metabolism regulation, were also low (p>0.05) testifying to certain activation of osteoclastogenesis (see Table 2).

## Conclusion

Our findings show that PTOA animals suffer rearrangement of skeletal tissues as compared to intact controls. This process involved the degenerative destruction of both cartilage and osseous tissues along with profound prooxidant activity and relative incompetence of the thiol system while the indicators of general antioxidant activity are within the normal range. These findings suggest the importance of analyzed makers for PTOA pathogenesis and define the prospects of their application for the evaluation of therapy response.

